# 3D microarchitecture of the human tuberculous granuloma

**DOI:** 10.1101/2020.06.14.149898

**Authors:** Gordon Wells, Joel N. Glasgow, Kievershen Nargan, Kapongo Lumamba, Rajhmun Madansein, Kameel Maharaj, Robert L. Hunter, Threnesan Naidoo, Pratista Ramdial, Llelani Coetzer, Stephan le Roux, Anton du Plessis, Adrie J.C. Steyn

## Abstract

Our current understanding of the pathophysiology of human pulmonary TB is limited by the paucity of human TB lung tissue for study and reliance on 2D analytical methods. Here, to overcome the limitations of conventional 2D histopathology, we used high-resolution 3D X-ray imaging (µCT/nCT) to characterize necrotic lesions within human tuberculous lung tissues in relation to the airways and vasculature. We observed marked heterogeneity in the 3D structure and volume of lesions. Also, 3D imaging of large human TB lung sections provides unanticipated new insight into the spatial organization of TB lesions in relation to airways and the vascular system. Contrary to the current dogma depicting granulomas as simple spherical structures, we show that TB lesions exhibit complex, cylindrical, branched-type morphologies, which are connected to, and shaped by, the small airways. Our results highlight the likelihood that a single structurally complex lesion could be wrongly viewed as multiple independent lesions when evaluated in 2D. These findings have strong implications for understanding the pathophysiology and evolution of TB disease and suggest that aerosolized drug delivery strategies for TB should be reconsidered.

## Introduction

Tuberculosis (TB) is a global infectious disease caused by the bacterium *Mycobacterium tuberculosis* (*Mtb*). Histopathological analysis was a mainstay for the investigation of TB disease throughout the 1940s^1^ and 1950s^2^ when post-mortem and resected human lung tissues were routinely available. These pioneering studies have strongly influenced our current understanding of the spectrum of tuberculous lesions, morphology and pathology.

The prevailing dogma in the TB field is that TB granulomas form spherical or ovoid structures within the parenchyma^3-11^. However, this assumption is not always supported by experimental evidence. Further, remarkably little is known about the structure of the caseous granuloma, the distinctive feature of infection by *Mtb* in humans. Hence, a deeper understanding of the human TB granuloma is urgently needed to more accurately inform preventive and therapeutic TB strategies.

The ability to examine pathological TB structures within large tissues in 3D could allow identification of disease-specific features and improve diagnosis. Hence, it is reasonable to speculate that the limitations of conventional histological analysis have begun to hinder more detailed examination of human TB pathophysiology in the current antibiotic era, especially with the emergence of HIV. Other factors contributing to our limited understanding are the reliance on animal models which do not recapitulate human pulmonary TB phenotypes, and the paucity of routinely available resected human TB lung tissues^12,13^. Therefore, imaging approaches that provide high-resolution digital 3D imaging of TB lesions will allow comprehensive analysis of the complex 3D microanatomical features specific to pulmonary TB.

X-ray computed tomography (CT) is an invaluable imaging tool for nondestructive assessment of tissue in medical diagnosis^14-16^. High resolution micro-CT (µCT) is typically used for materials with high electron density and lends itself to *ex vivo* analysis of pathologies involving bone structure or calcium deposition^17^. Imaging of soft tissue can be improved by addition of electron-dense contrast agents (*e*.*g*., iodine, osmium, or tungsten) or using high-energy flux monochromatic x-rays generated by synchrotrons. To our knowledge, however, no study has reported the use of µCT to examine bacterial or viral disease in human lungs. While µCT is an experimental imaging modality, high-resolution computed X-ray tomography (HRCT) is often used clinically to aid TB diagnosis^18,19^. Specifically, HRCT can detect phenotypes of *Mtb* infection such as bronchiectasis, cavity formation and tissue consolidation^20^. While HRCT is non-invasive, it suffers from lower resolution (∼0.23-1.5 mm) and usually requires a contrast agent for imaging of homogeneous soft tissue^21,22^.

Here, we characterized the 3D environment of the human tuberculous lung *ex vivo*. We examined the 3D structure of TB granulomas, their spatial position relative to the airways and vasculature, and confirmed our findings using histopathology and immunohistochemistry. Overall, we demonstrate the utility of µCT for direct visualization of pulmonary TB in detail, thereby advancing our understanding of how *Mtb* causes destructive human TB.

## Results

### µCT characterization of the human TB granuloma

In Durban, South Africa, *Mtb*-infected human lung tissues are routinely obtained following resection of irreversibly damaged lung regions exhibiting bronchiectasis and/or cavitary lung disease^23,24^. For this study, we analyzed formalin fixed (FF) lung specimens obtained from 17 subjects (Table 1). Comprehensive sampling from different regions of each lung or lobe allowed us to evaluate microenvironments at different stages of tissue pathology. Gross architectural distortion with conspicuous upper lobe cavitation in a background of bronchiectasis, lung shrinkage, and fibrosis were noted in cut sections of several specimens. The specimens exhibited typical features of bronchiectasis and contained tubercles of varying size and shape. Representative sections of the cavitational and parenchymal abnormalities were used for imaging studies; see Table 1.

**Table 1.** Clinical characteristics of human subjects.

| # | Patient # | Age | Sex | Type | Macroscopic and Microscopic features | Type of resection |
| --- | --- | --- | --- | --- | --- | --- |
| 1 | 202134 | 38 | F | TB | Caseative necrosis, 2 cm cavity in upper lobe, identifiable acid-fast bacilli, suppurative granulomas with microabscess formation | Left pneumonectomy |
| 2 | 202203 | 67 | M | TB | Necrotizing granulomatous inflammation with acid-fast bacilli, intra-alveolar foamy macrophages are present | Left upper lobe lobectomy, |
| 3 | 202211 | 32 | F | TB | Cavitation and miliary TB, fibrocalcific nodules, acid-fast bacilli present within macrophages, features of lymphoid interstitial pneumonia are present | Left pneumonectomy |
| 4 | 202294 | 39 | F | TB | Necrotizing granulomatous inflammation and hemorrhage, granulomas with central microabscesses are present, acid-fast bacilli stain is positive | Left upper lobe lobectomy |
| 5 | 202254 | 37 | F | TB | Caseous necrosis with cavity containing necrotic material, necrotizing granulomatous inflammation, Langhan's giant cells are present, paucibacillary | Right lower lobe lobectomy |
| 6 | 202733 | 37 | F | TB | Multiple round tubercles were noted, necrotizing and non-necrotizing granulomatous inflammation, acid-fast bacilli positive, Langhans giant cells are present, alveolar spaces filled with eosinophilic material and hemorrhage | Right upper lobe lobectomy |
| 7 | 202194 | 15 | F | TB | Multiple tubercles confirm active necrotizing granulomas, acid-fast bacilli were seen, vasculitis, interstitial fibrosis | Right upper lobe lobectomy |
| 8 | 202172 | 28 | M | TB | Shrunken, collapsed tubercles appear irregular in shape, calcified foci were confirmed, focal tuberculous scars and fibrocaseous foci were noted, intra-alveolar siderophages, paucibacillary | Left upper lobe lobectomy |
| 9 | 202156 | 33 | F | TB | Shrunken lung, collapsed with bronchiectasis and fibrosis, tubercles irregular in shape, irregular cavity was noted, granulation tissue is prominent, aspergillosis, paucibacillary | Left pneumonectomy |
| 10 | 202187 | 45 | F | XDR TB | Necrotizing granulomatous inflammation, granulomas surrounding central areas of caseous necrosis, Langhan's giant cells are present, cavities contained neutrophils, acid-fast bacilli are present | Right pneumonectomy |
| 11 | 202510 | 50 | M | TB | Bronchiectatic, contains cavity wall lined by chronic suppurative inflammation, granulation tissue and foamy histiocytes were observed, aspergilloma was present, hemorrhage, or oedema and organizing pneumonia were seen | Right pneumonectomy |
| 12 | 202126 | 37 | F | TB | Bronchiectatic cavitation and caseous necrosis, fibrocalcific and fibrocaseous granulomas were noted, histiocytes, giant cells and central areas of necrosis predominate, intra-alveolar foamy histocytes, paucibacillary | Right pneumonectomy |
| 13 | 202340 | 65 | F | TB | Irregular tubercles with bronchopneumonia were identified, extensive granulomatous inflammation, acid-fast bacilli were identified, interstitial fibrosis and areas of gross scarring | Right pneumonectomy |
| 14 | 202113 | 24 | M | TB | Necrotizing granulomatous inflammation with acid-fast bacilli, some of the granulomas contained suppurative centres, acid-fast bacilli were identified, granulomatous process is also seen within the hilar lymph nodes | Left upper lobe lobectomy |
| 15 | 202221h |  |  |  | Healthy lung control |  |
| 16 | 202254h |  |  |  | Healthy lung control |  |
| 17 | 202223h |  |  |  | Healthy lung control |  |

To improve the clinicopathological analysis of TB, we attempted to establish a correspondence between X-ray density and pathological features within lesions that permit 3D reconstruction. TB lesions have pathological features that can evolve over decades^1,2^. While these structures likely represent unique immunopathological microenvironments, their contribution to TB disease and persistence of *Mtb* is poorly understood. This partly due to the inability of 2D histology to adequately characterize these deformities. We scanned a contrast-stained lung tissue sample (15 × 10 × 10 mm) with caseous necrosis (Figure 1A) at 12.0 µm resolution (Figure 1B). Segmentation identified distinct regions that matched blood vessels and necrotic lesions. The identification of lesions and vasculature with µCT was confirmed by histology using H&E (Figure 1C) and trichrome staining (Figure 1D), revealing several necrotic lesions and evidence of fibrosis. Inspection of the lesions revealed common electron density features, which we confirmed quantitatively by plotting relative X-ray attenuation (electron density) across representative sections (Figure 1E, F). First, the necrotic lesions are surrounded by a dark outer layer of CT intensity (Figure 1G-J) corresponding to lamellar fibrosis by H&E (Figure 1K-N) and trichrome staining (Figure 1O-R). Second, the necrotic region itself exhibits a light border (Figure 1G-J) that corresponds to a more intensely stained border in H&E and trichrome staining (Figure 1K-R), surrounding a mass of less electron-dense necrotic material (Figure 1G-J). Hence, we were able to establish a correlation between pathophysiological features and changes in electron density. Additionally, in one lesion (Figure 1G), µCT revealed two internal “lobe-like” lesions that are not apparent in the corresponding histopathology (Figure 1K, O), further emphasizing the potential of µCT to identify unusual pathological features.

**Figure 1.**
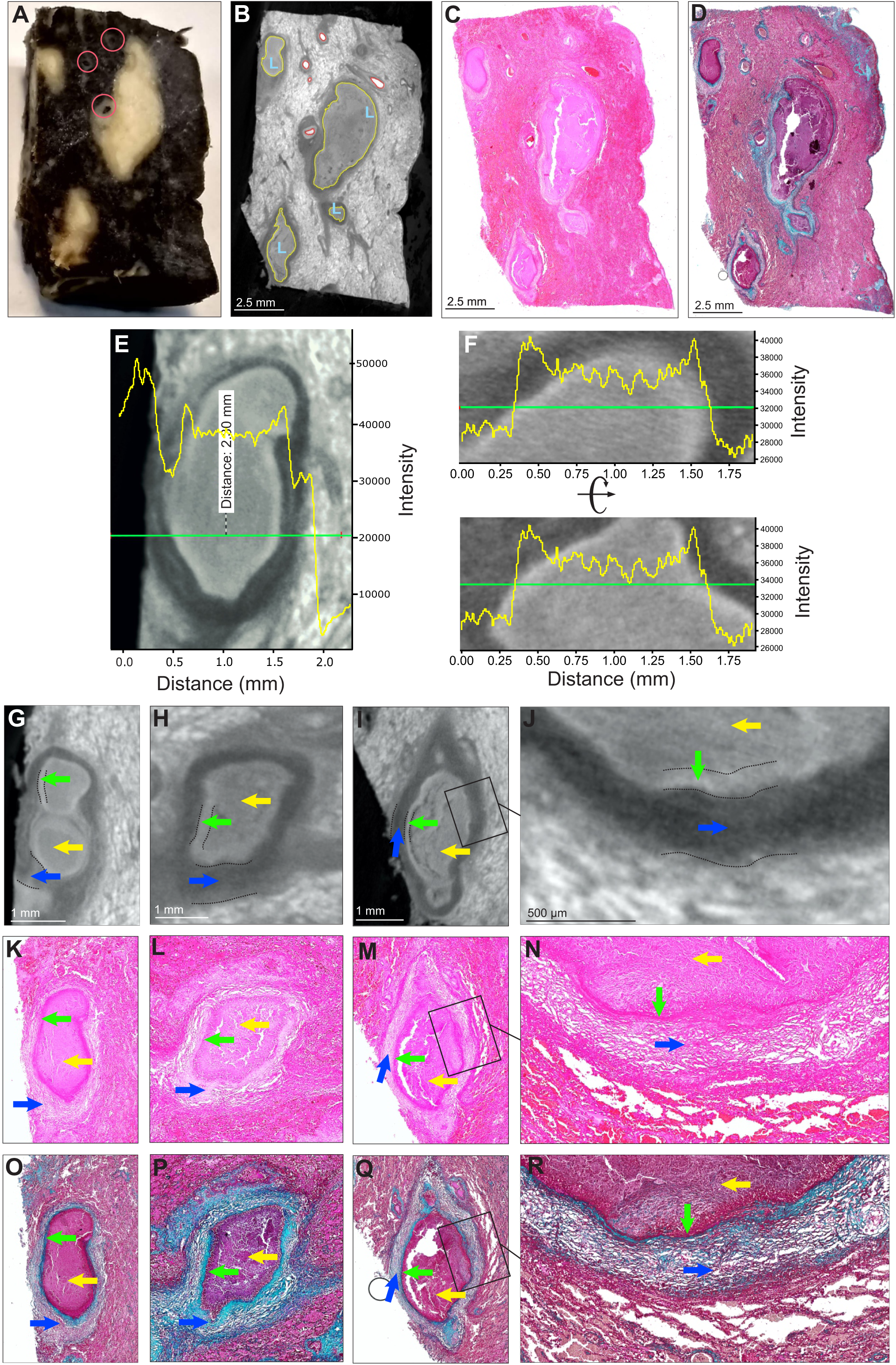
µCT and histology of human TB-lung with caseous necrosis (Sample A) **(A)** Gross image of sample E exhibiting caseous necrosis. Pink circles indicate blood vessels. (**B)** µCT (12.0 µm resolution) of sample E, caseous necrosis (yellow, “L”), hemorrhage/blood (red). (**C**) H&E histology of (B) (**D**) MT histology of (B). (**E-J**) Necrotic regions have a ‘halo-like’ appearance, with a slightly brighter outer shell (green arrows) surrounding a slightly darker interior (yellow arrows). Necrotic regions are surrounded by a dark border (blue arrows). (**E, F**) Typical X-ray opacity profile/electron density (yellow graph) across necrotic lesions measured along the green axis. Dark fibrotic regions are followed by a slightly opaquer ring that surrounds the (lighter) lesion. (**G-J**) Representative µCT images of caseous necrotic lesions. (**K-N**) H&E and (**O-R**) MT histology corresponding to panels G-J reveal the darker shell (blue arrows) surrounding the necrotic regions that corresponds to fibrotic tissue.

Overall, these data demonstrate that µCT can effectively complement standard histopathological analysis by revealing hidden pathological features that might otherwise be disregarded by pathologists.

### 3D segmentation and spatial distribution of TB lesions in the human lung

While conventional histopathological analysis provides detailed information on very small areas of interest, it cannot contextualize TB lesions within the overall lung architecture. This has limited our understanding of the distribution and shapes of lesions within the human TB lung. µCT has the potential to improve our understanding of the evolution of granuloma formation and structure, relative to the diffusion of drugs, O_2_ and nutrients. To contextualize TB lesions relative to the vasculature and airways, we used µCT to scan a large slice (∼14 × 1.5 × 6 cm) of infected lung at 52.0 µm resolution (Figure 2A, Figure S1, Video S1). 3D segmentation shows that larger lesions are oriented with a directionality similar to the vasculature and bronchial networks (Figure 2B). Notably, as is evident by the multiple lesions that were curtailed during sectioning of the tissue (Figure 2B, lower image), the data suggest that the lesions are more complex as they continue beyond the sectioning plane. Additionally, airways are absent in areas where lesions predominate, suggesting that lesion inception through bronchial obstruction has replaced the former airways.

**Figure 2.**
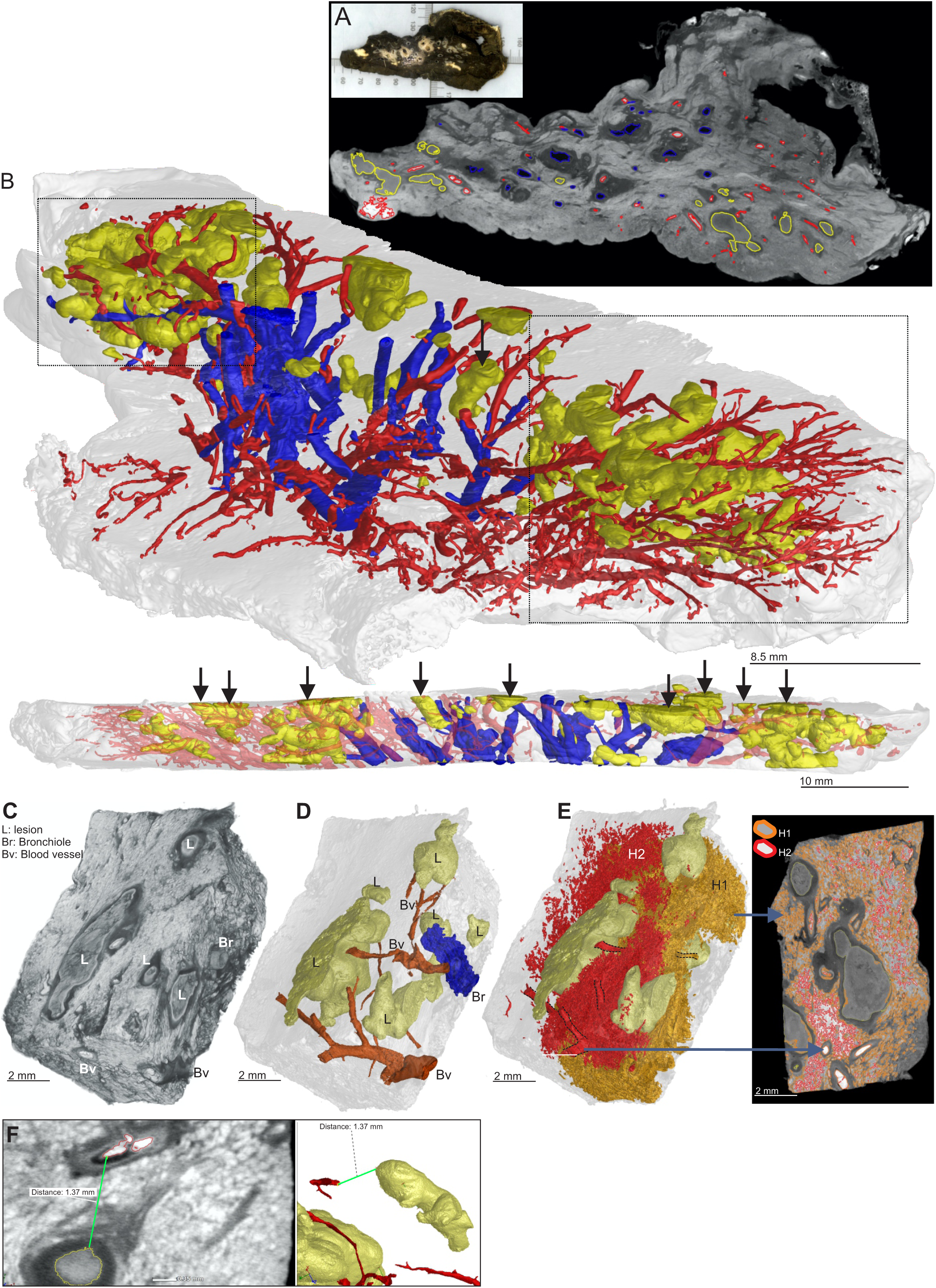
µCT and segmentation of human TB-lung with caseous necrosis (Sample B) (**A**) 2D slice of µCT (52.0 µm resolution) of a human lung lobe. Necrotic lesions (yellow), bronchi/bronchioles (blue) and vasculature (red) are outlined. (**B**) Complex necrotic lesions are oriented similarly to the airways and vasculature. 3D renderings of lesions (yellow), bronchi/bronchioles (blue) and vasculature (red) segmentation. Lower image: side view of **A** with truncated lesions (also observed in **D**) indicated by vertical arrows. (**C**) Sample B exhibiting caseous necrosis. ScatterHQ (VGL Studio) rendering of surface electron density. L, truncated lesion; Br, bronchiole/airway; Bv, blood vessel. (**D**) 3D segmentation of blood vessels (red, Bv), airways (blue, Br) and necrotic lesions (yellow, L). (**E**) 3D segmentation of lesions (yellow) and hemorrhage (red/orange). Blood vessels and nearby regions of bleeding stain brightly, with decreasing intensity further away from blood vessels. By selecting all regions above a high intensity threshold (red, H2), hemorrhaging (including intact vasculature) can be quickly segmented. The lower threshold (orange, H1) also selects other components outside hemorrhaged region (*e*.*g*, within the lesions). (**F**) Representative µCT slice of segmented regions from (**D**) demonstrating the distance between the lesions and the vasculature.

Surface area rendering of a sub-section of this sample distinctly identified lesions, blood vessels and airways (Figure 2C). 3D segmentation revealed six lesions (Figure 2D) with volumes of 23.89, 6.30, 4.09, 4.00, 0.56, and 0.49 mm^3^ (total: 39.33 mm^3^, 6.9 % of total tissue volume). Intense staining of erythrocytes permitted a rapid, albeit partial reconstruction of the vasculature (Figure 2E). Vascular destruction, also observed in Figure 2B, contributes to interstitial hemorrhage resulting in nutrient and O_2_ deprivation which further contribute to TB disease. A considerable degree of hemorrhage was observed with segmentation by thresholding, generating large complex regions of interest obscuring the lesions and airways (Figure 2E). There was little, if any, healthy functional lung tissue within this sample. Lastly, we measured distances between intact vasculature and necrotic lesions. This proximity would almost certainly impact lesion development and morphology. The maximum O_2_ diffusion distance is 100-200 µm from a blood vessel^25-27^, and metabolic zonation may account for spatial lesion heterogeneities^28^. Although histopathological analyses have shown that TB lesion distance from the vasculature can exceed 200 µm, this is not conclusive and could be influenced by the sectioning plane. Using 3D segmentation, we observed that geometrically, the vasculature follows the curvature of the lesions. The distances between blood vessels and lesions range from ∼0.5 - 1.4 mm (Figures 2F, S2, S3A, S3B). Hence, the curvature of lesions must be considered in order to accurately measure these distances, which are a crucial factor in understanding how the vasculature delivers nutrients, drugs and O_2_ to bacilli (Figure S3B) and immune cells in and around the lesion.

Our results show that integration of conventional 2D histopathological methods with µCT provides the means to identify key pathological features such as lesion volume, 3D structure, and intralesional features in the context of the whole lung. The spatial organization of lesions proximal to the pulmonary vasculature is particularly important, since vascular destruction will reduce delivery of anti-TB drugs, O_2_ and nutrients. The lack of airways and the directionality of lesions that accords with the vasculature suggest that TB lesions observed by conventional histopathology may sometimes be cross-sections of obstructed airways instead of spherical lesions. Hence, application of µCT has substantial potential to advance our understanding of the pathophysiological mechanisms of TB disease and poor response to anti-TB drugs.

### Complex 3D structures of TB lesions and communication with the airways

Here, we characterized the spectrum of caseous lesion structures (Figure 3A) obtained from the sample in Figure 2A, B in more detail. In contrast to the current dogma that TB lesions are near-spherical or ovoid, segmentation of caseous necrotic lesions revealed remarkable morphological heterogeneity and complexity (Figure 2A). Of the 40 lesions segmented in Figure 2A, B, multiple highly branched structures were observed, in contrast to the expected ovoid form. The radius (of the smallest enclosing sphere) of these lesions ranged from 0.5-7 mm for the more elaborate forms. While smaller lesions where more spherical, larger lesions were branched with lower sphericity, which ranged from 0.23-0.6 for all lesions (1.0 is a perfect sphere) (Figure 3B).

**Figure 3.**
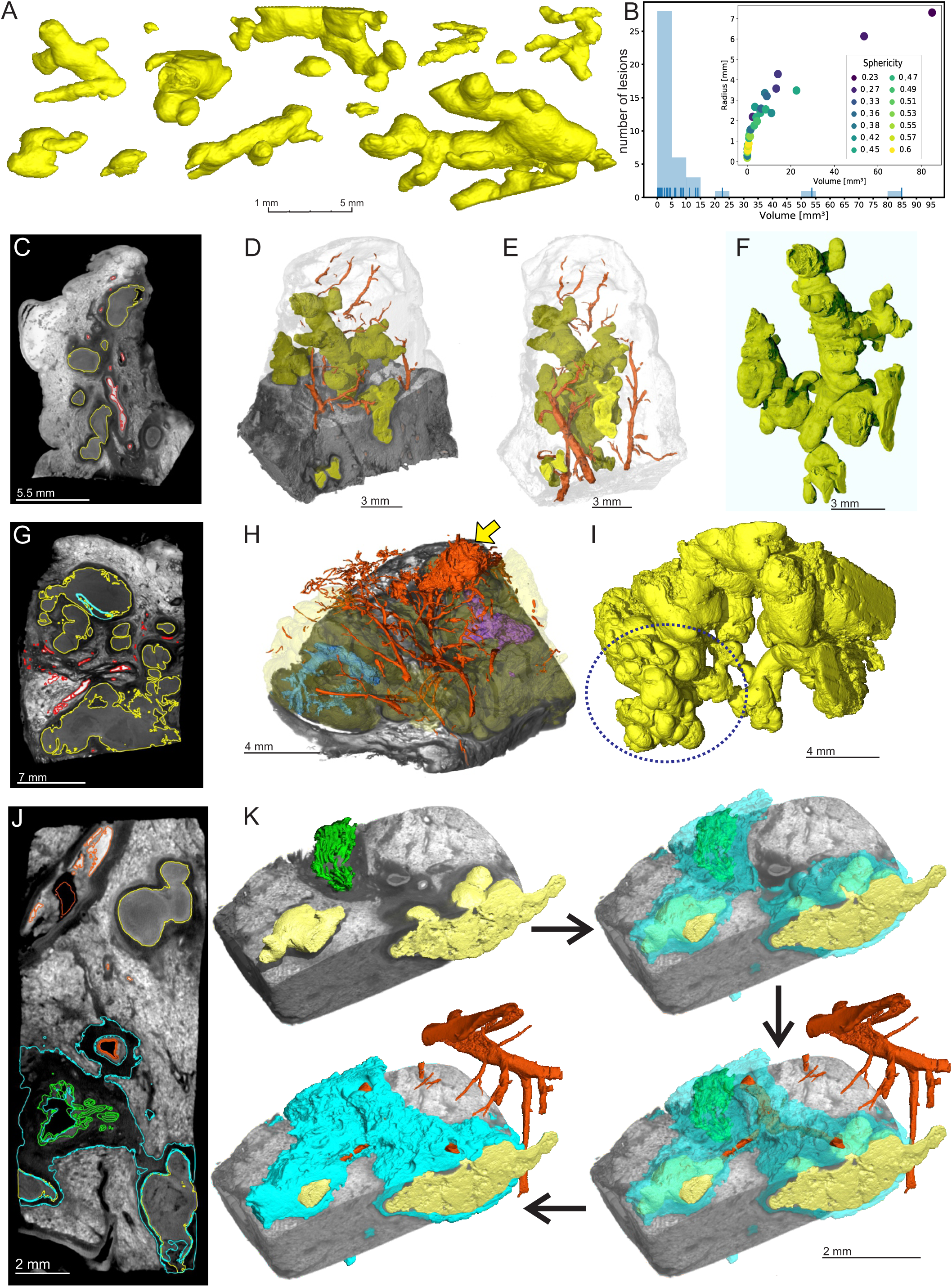
Lesion morphological heterogeneity and micro-structure of lesions and surrounding vasculature (Samples A-D) (**A**) The morphology of necrotic lesions in advanced TB (Sample B) ranges from small nodules (mm scale) to large branched structures (cm scale) within a lung sample. (**B**) Relationship between lesion size and shape in Sample B. Sphericity is the ratio of the surface area of the sphere with the same lesion volume to the lesion surface area. Smaller lesions tend to be nodular (higher sphericity), while larger lesions exhibit more complex shapes with a lower sphericity. (**C-F**) µCT of Sample C, obtained from the tip of Sample B. (**C**) 2D slice of tip of Sample B. (**D**) 3D rendering of lesion (yellow) and vasculature (red) segmentation in relation to X-ray absorption/electron density. (**E**) 3D rendering of lesion (yellow) and vasculature (red) segmentation in relation to sample surface. (**F**) 3D rendering of lesion (yellow) only. (**G-I**) µCT of Sample D. (**G**) 2D slice of Sample D. (**H**) 3D rendering of lesion (yellow) and vasculature (red) segmentation in relation to X-ray absorption/electron density. Yellow arrow; hemorrhage, purple and turquoise structures; obliterated airways. (**I**) 3D rendering of lesion (yellow) only. Dotted circle: area resembling tree-in-bud. (**J**) 2D slice of Sample A (excised from Sample B). (**K**) 3D rendering of lesions (yellow), vasculature (red), bronchus (green) and lesions/vasculature/airway connections (cyan) from (**J**).

Smaller lung samples with caseous necrosis were excised and scanned at higher resolution, further revealing the complexity of the lesion microenvironment (Figure 3C-K, Video S2). One section taken from the tip of the sample in Figure 2A, B contained a complex ginger root-like structure (Figures 3C-F, Video S3). The sample in Video S3 and a second sample from a different patient revealed small lobular regions resembling the buds of the “tree-in-bud” signature often seen in HRCT scans^29,30^ of severe TB (Figures 3H, I, Video S4). We observed severe hemorrhage as is indicated by the white contrast areas in Figures 3C, G and J and vascular destruction in Figure 3H, as well as intricate vasculature that surrounds lesions in both samples (Figures 3D, E, H). Both the structures in Figures 3C-F and G-I continue beyond the scanned view, indicating that the native, uncut lesions were larger and likely more complex.

To further explore the connection between bronchi and TB lesions, i.e., obstructed airways, we segmented the volumes surrounding lesions, airways and vasculature shown in Figure 2A, B. µCT reveals darker regions of similar radio-opacity surrounding the necrotic lesions, airways and blood vessels (Figure 3J). Segmentation of this volume (indicated in cyan) reveals that it surrounds and connects granulomas with airways and blood vessels (Figure 3K). This further suggests that the shape of TB lesions is dictated by the small airways (Video S5).

Overall, we demonstrate that TB granulomas are remarkably structurally diverse and have multifaceted connections with the surrounding vasculature and airways. Although 2D histopathology sectioning typically reveals “round” granulomas that are intuitively inferred to be spherical, our findings challenge this prevailing dogma. Rather, our results point to a more complex, cylindrical or branched-type morphology for advanced TB lesions, which are connected and shaped by the small airways.

### TB lesion formation through bronchial obstruction

Our µCT data suggest that TB granuloma structure is influenced by the small airways. Here, we confirm this finding by examining immune cell infiltration and subsequent blockage of small airways using immunohistochemistry (IHC). Firstly, we confirmed that *Mtb* bacilli were present intracellularly in macrophages and neutrophils (Figures S4, S5), extracellularly within alveoli (Figures S4, S5) and within an obstructed bronchus (Figure S6). Next, in highly consolidated areas of the tuberculous lung (Figure 4A, B and Figure S7), we identified patterns of epithelial cell remnants consistent with obstructed small airways, as indicated by cytokeratin 7 (CK7), and 3-mercaptopyruvate sulfur-transferase (3MPST) staining, which is specific for epithelial cells^31^ (Figure 4C-F). This finding demonstrates that immune cell recruitment during TB inflammation can obstruct the small airways, which can further develop into a granuloma surrounded by epithelial cells (Figure S8). In less consolidated areas, macrophages, neutrophils and lymphocytes obstruct alveoli (Figure 4G) leading to independent and coalesced granulomas (Figure 4H), indicative of the early stages of alveolar consolidation.

**Figure 4.**
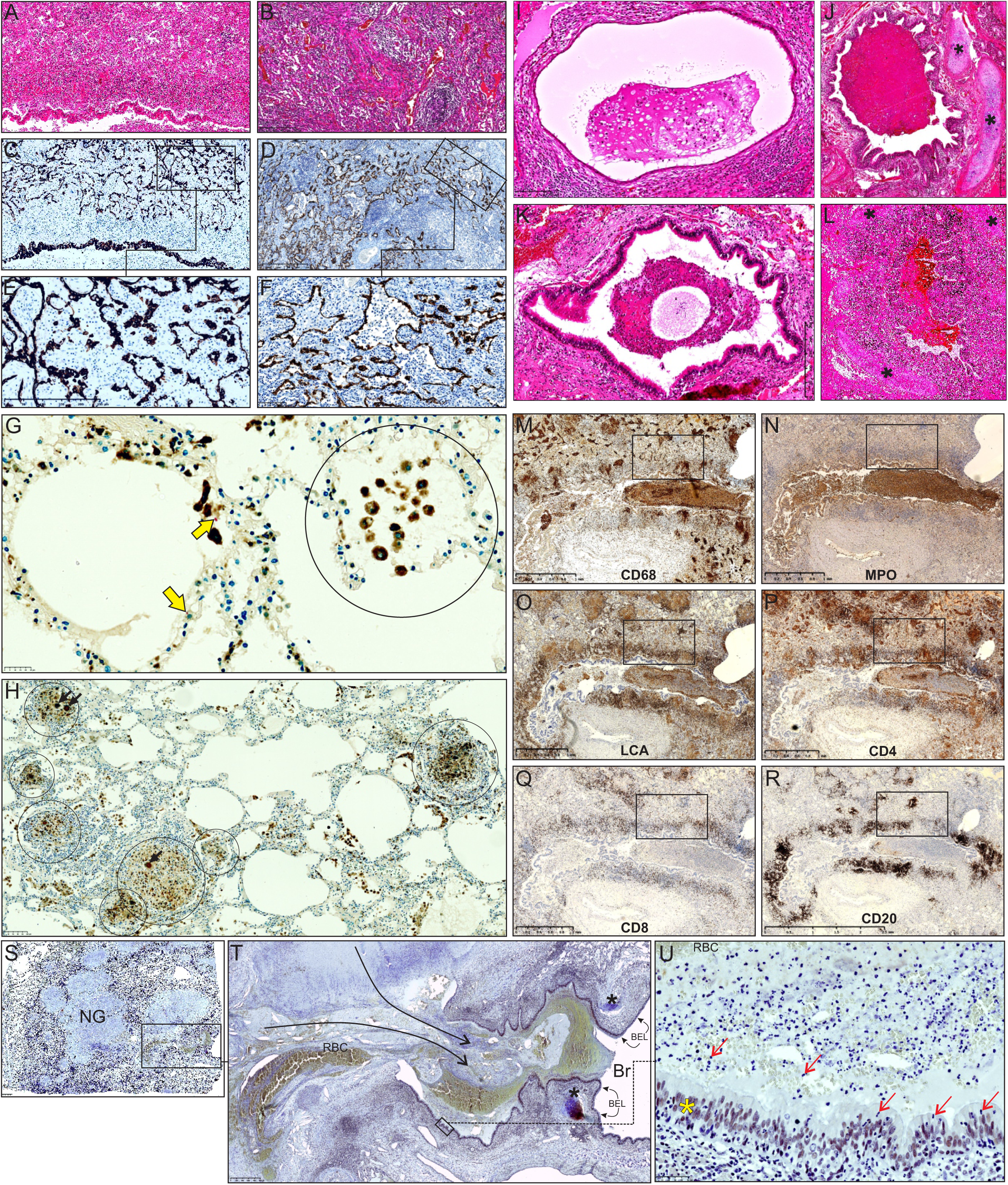
Histopathology of the small airways of an *Mtb*-infected human lung. (**A, B**) Low power magnification of H&E stain in lung parenchyma. (**C, D**) Low power and (**E, F**) medium power magnification of epithelial staining in the adluminal layer (**C, E**; CK7, **D, F**; 3MPST). (**G, H**) Combined CD68 and ZN staining. Circled areas: alveoli filled with macrophages, arrows; giant cells. Yellow arrows; *Mtb*. (**I, J, K, L**) H&E staining of bronchial obstruction. (**M, N, O, P, Q, R**) IHC of myeloid and lymphocytes. Boxed areas; see Figures S7 and S9. (**S**) Low power magnification of ASM (nuclear) staining. Note the spilling of necrotic material from granulomas (NG) into an airway. (**T**) Medium power image (black asterisks, cartilage, BEL; bronchial epithelial layer, black arrows; spillage of necrotic material into bronchus, Br; bronchus). (**U**) High power depiction of the BEL (yellow asterisk) with immune cells in the airway. Red arrows; neutrophils. RBC; red blood cells. See Figure S10 for high-power image.

Next, we examined the contribution of innate and adaptive immune cells to bronchial and alveolar obstruction. Histopathological appraisal of lung tissue specimens from several TB patients identified numerous obstructed bronchi containing immune cells (Figure 4I-L). We identified an abundance of myeloid cell populations, indicated by strong positive staining of leukocyte common antigen (LCA), myeloid peroxidase (MPO), and CD68 in these cells (Figure 4M, N, O; see Figure S7 for higher power image). Positive staining of CD4+ and CD8+ lymphocytes inside and outside the obstructed bronchus (Figure 4P, Q), and CD20+ cells (Figure 4R) that dominate the area around the bronchus, was observed. Notably, in the consolidated diseased areas in Figure 4M-R (boxes), we observed clear evidence of myeloid cell and lymphoid cell infiltration by IHC (Figure S9A-F), which is in support of the consolidation shown in Figure 4A-F. Lastly, we observed necrotic material and immune cells from TB granulomas spilling into a bronchus (Figure 4S-U; see Figure S10 for high power image), providing compelling evidence for expansion of necrotic lesions along the airway network to help shape granuloma structure. These granulomas are surrounded by foam cells (Figure S11), consistent with historical studies showing that obstructive lobular pneumonia softens lung tissue (i.e., caseating necrosis), which is then coughed up, leading to cavitation^13,32^.

In conclusion, histopathology and IHC data are fully consistent with our µ/nCT data demonstrating that recruitment and expansion of immune cells in the airways, eventually followed by necrosis, contribute to blockage of the airways and the 3D shape of the granuloma. These findings have implications for how TB transmission is triggered through coughing, for cavity formation, and for aerosolized drug delivery strategies.

## Discussion

While conventional histological methods have been the gold standard for appraising TB disease pathology for over 100 years, there is a need to address the multidimensionality of diseased tissue using advanced high-resolution imaging modalities. Our current understanding of TB disease has been shaped by the histopathological interpretations in the 1940s and 1950s by Arnold R. Rich^1^, George Canetti^2^ and Edgar M. Medlar^33,34^ when post-mortem and resected human lung tissues were routinely available. With the emergence of HIV and its synergism with TB, and broad access to anti-TB drugs, all of which influence disease pathology^35-37^, it is reasonable to suppose that TB pathology phenotypes have changed. Difficulty in describing human pulmonary TB disease has hampered the TB field for decades, whereas the use of animal models for TB has flourished. Unfortunately, no single animal model accurately duplicates the full spectrum of human pulmonary TB phenotypes. Here, µCT imaging has provided new insight into the morphologies of human necrotic TB lesions, demonstrating that they can form branched and cylindrical structures with large variations in volume, size and spatial distribution and that they are connected to the small airways. These findings contrast with the current dogma that granulomas are spherical, an understandable conclusion based on conventional histopathology. Our findings exemplify how 3D visualization of TB disease pathophysiology can improve our understanding of the evolution of TB granuloma and provide a foundation for a 3D atlas of the human tuberculous lung. Lastly, our findings establish a clinically relevant framework for the discovery of imaging biomarkers as diagnostic indicators, and provide a strong rationale for development of aerosolized anti-TB drug delivery strategies.

A significant advance in this study is the application of 3D segmentation to the microarchitecture of the tuberculous lung, which provides detailed insight into the spatial relationship between TB granulomas, airways, and the vascular system. To our knowledge, such findings have not yet been reported for any pulmonary pathogen, bacterial or viral. Several unexpected discoveries about the TB granuloma were made. Firstly, demonstrating that the TB granuloma represents a spectrum of complex, branched-type morphologies, and is shaped by the small airways, has implications for understanding the evolution of granuloma, of which little is known. This new insight represents an important advance with strong clinical implications since the prevailing presumption has been that the granuloma is spherical^3-11^. Also, our 3D segmentation highlights the possibility that a single structurally complex lesion could be erroneously viewed as multiple independent lesions when evaluated in 2D. The potential for misinterpretation of granuloma number, size or position indicates that great care must be taken while interpreting “-omic” data derived directly from TB lesions, as conclusions will be influenced by the actual (but unknown) 3D shape of the lesion. Further, conclusions drawn regarding microenvironments surrounding what appear to be multiple granulomas could change if it were understood that a single complex lesion was under investigation. Secondly, our findings highlight the pathophysiological factors that help dictate the shape of the granuloma. Here, it is evident that immune cell infiltration in the alveoli, bronchioles and bronchi dictate the shape, and that immune cell recruitment and subsequent necrosis expand in the airways to follow “the path of least resistance”. Alveolar walls contain numerous inter-alveolar pores that may function as conduits for the dissemination of *Mtb* or infected cells. Also, several granulomas in this study are reminiscent of the tree-in-bud form, an HRCT signature that is present in virtually all cases of active pulmonary TB^29,30^. Thirdly, the spatial relationship between TB lesions and pruning of the surrounding vasculature, which impedes the delivery of nutrients, O_2_, anti-TB drugs and immune cells to granulomas, may shed light on the complex pathophysiology involved in promoting persistence and drug tolerance. For example, 3D renderings of the vascular system from diseased TB lungs show destruction of the vascular network, which would reduce delivery of anti-TB drugs, metabolites, and O_2_. This may explain why drugs do not reach bactericidal concentrations within TB lesions^38^ and how *Mtb*, which requires O_2_ to grow, is able to persist long-term in O_2_-deficient lesions, presumably in a state of metabolic shutdown. For instance, the maximum O_2_ diffusion distance is ∼200 µm^25-27^ after which tissue becomes hypoxic. By accurately measuring the distance between blood vessels and the segmented TB lesions, we conclude that many necrotic lesions are hypoxic. Based on a recent study^28^, it is almost certain that gradients, or zones, of drugs, metabolites and O_2_ exist within TB lung tissue. Therefore, separate anisotropic gradients for different drugs^38,39^ may trigger sequential development of *Mtb* drug resistance or tolerance by passaging through environments with low drug concentrations. Therefore, therapeutic angiogenesis and aerosolized drug delivery strategies^40^ may represent plausible approaches to increase anti-TB drug levels in the granuloma.

Our findings also suggest our approach is transformative for histopathological assessment as it will contribute to a more informative clinicopathological analysis for TB. Notably, all our histological sections could be matched to the corresponding sliced plane from the µCT 3D volume. Consistent with autopsy studies^20^, these findings provide further insight into the evolution of TB lesions, and suggest that necrotic material fills the bronchiolar lumen to induce bronchial wall necrosis, which promotes progressive necrosis of the lesion. Furthermore, integration of µCT imaging with histopathology has strong potential to influence other disciplines including pathology, biomedical imaging, infectious diseases and cancer, ultimately leading to new discoveries. Overall, our data demonstrate that µ/nCT is a powerful imaging tool to study the mechanism of granuloma formation.

Our study has some limitations. First, this was a focused study and a limited number of lung tissue samples were examined from TB patients with diverse medical histories and treatments; hence it is likely that a larger test cohort may render a more representative disease spectrum. However, sampling from different regions of each lung allowed us to evaluate microenvironments at different stages of tissue pathology. Second, we used iodine, a widely employed contrast agent, in our studies; however, other agents may provide unique staining patterns that allow identification of different adjacent tissues. This is especially true if reversible staining protocols can be developed that allow serial staining with different contrast agents. Contrast staining with iodine also interfered with subsequent hematoxylin staining for histological follow-up and requires further optimization. Lastly, similar to conventional histopathology, shrinkage of tissue during formalin fixation and staining is widely known and may influence volume calculations. However, this could be mitigated by use of polyoxometalates^41^.

Overall, our findings have important implications for TB disease treatment and diagnosis as several surprising findings were made, including the spectrum of granuloma 3D structures, the size and volume of TB lesions, and their spatial organization in relation to the vasculature and airways. Secondly, scouting for pathological features may help guide and expedite histopathological follow-up studies. Thirdly, digitized 3D image libraries of tissue and organs from TB patients could be used to identify novel imaging biomarkers based on patterns of differential radio-opacities^42^ and establishment of a 3D reference atlas of the tuberculous lung. Lastly, our findings suggest that aerosolized anti-TB drug delivery strategies for the control of TB should be reconsidered.

## Methods

### Ethics and Human Subjects

This study was approved by the University of KwaZulu-Natal Biomedical Research Ethics Committee (Class approval study number BCA 535/16). Patients undergoing lung resection for TB (Study ID: BE 019/13) were recruited from King DinuZulu Hospital Complex, a tertiary center for TB patients in Durban, South Africa. *Mtb*-infected human lung tissues are routinely obtained following surgery for removal of irreversibly damaged lobes or lungs (bronchiectasis and/or cavitary lung disease). Written informed consent was obtained from all participants. All patients undergoing lung resection for TB had completed a full 6-9-month course of anti-TB treatment, or up to 2 years of treatment for drug-resistant TB. Patients were assessed for extent of pulmonary disease (cavitation and or bronchiectasis) via HRCT. The fitness of each patient to withstand a thoracotomy and lung resection was determined by Karnofsky score, six-minute walk test, spirometry and arterial blood gas. Assessment of patients with massive hemoptysis included their general condition, effort tolerance prior to hemoptysis, arterial blood gas measurement, serum albumin level and HRCT imaging of the chest. On gross assessment, all pneumonectomies or lobectomies were bronchiectatic, hemorrhagic, variably fibrotic and atelectatic and contained visible tubercles (Table 1).

### Sample Preparation

Seven tissue samples (Samples A – G, Table 2) from resected human lungs (un-inflated) (Table 1) were selected for µ/nCT analyses. All samples were fixed in 10% buffered formalin for at least 14 days. Samples A and B were obtained from resected lungs with evidence of cavitation and *Aspergillus* infection in sample B. Samples C and D represent relatively healthy tissue from a cancerous lung and *Mtb*-infected lung, respectively. Sample E was selected from a lung with evidence of severe TB infection including extensive caseous necrosis. Samples F and G exhibit calcification, as well as fungal infection in F. Samples B-E were contrast stained with iodine by immersing the samples in 2.5 % Lugol’s solution for 1-5 days depending on the size of the sample. Samples F and G were also mounted in paraffin wax blocks before scanning.

**Table 2.** Scanning settings. All µCT scans were performed at 16-bit depth

| Figures | Sample | Target | Instrument | Scan time (s) | Averaging | Skip | Resolution (μm) | Voltage (kV) | Current (μA) | Images | Preparation |
| --- | --- | --- | --- | --- | --- | --- | --- | --- | --- | --- | --- |
| Figure 1B, E-J<br>Figure 2C-F<br>Figure 3J, K.<br>Figure S9<br>Video S5 | A | W | μCT | 3000 | 5 | 1 | 12.0 | 160 | 200 | 3000 | Contrast stained with I<sub>2</sub> |
| Figure 6A<br>Video S1 | B | W | μCT | 3000 | 5 | 1 | 52.0 | 160 | 260 | 3000 | Contrast stained with I<sub>2</sub> |
| Figure 2A-F<br>Video S2, S3 | C | W | μCT | 2700 | 5 | 1 | 15.0 | 160 | 180 | 2700 | |
| Figure 2G-I<br>Figure S8<br>Video S4 | D | W | μCT | 2800 | 5 | 1 | 15.0 | 80 | 400 | 2800 | Contrast stained with I<sub>2</sub> |

For µ/n-CT scanning, samples were mounted on or in 50 ml falcon tubes using a combination of cellophane tape and florist foam. Non-paraffin embedded samples were lodged above ∼ 5 ml formalin in the bottom of the tube with polystyrene foam and lodged between the walls of the tube to prevent shifting of the sample. The low density of polystyrene foam also enables easy deletion from the reconstructed volume during subsequent visualization and analysis. The tube was then sealed with parafilm for the duration of the scan to maintain a moist atmosphere and prevent desiccation. Prior to mounting, samples were rinsed with water and dabbed dry to remove excess staining solution.

### µ/nCT scanning

A General Electric Phoenix V|TomeX L240 system was used for µCT (2024×2024 pixel image, 16 bit depth) was used for µCT with a resolution range of 12.0 – 60.0 µm. A General Electric Phoenix Nanotom S (2304×2304 pixel image, 16 bit depth) was used for nCT with a resolution (isotropic voxel size) range of 4.1 - 16.0 µm. Although the instrument is capable of sub-µm resolution for small samples, none of the samples analyzed in this study were small enough. All samples were scanned over 360°. A range of settings were used to scan the samples as described further in Table 1. Briefly, voltage varied between 50-160 kV, current varied between 200-1000 µA and scanning times ranged from 2000-5400 seconds. For most scans a tungsten target was employed. A molybdenum target was used for two n-CT scans (Table 2).

### Image Processing and volume rendering

Volumes were reconstructed with system-supplied General Electric Datos software. Subsequent visualization and analysis (such as volume and density calculations) were performed in Volume Graphics VGStudio Max 3.1 or 3.2. Where possible, simple thresholding was employed for segmentation (demarcation of 3D regions of interest), followed by semi-automated segmentation using the VGStudio Max region growing tool. The region growing tool allows for manual selection of a 3D scan region based on adjustable intensity thresholds and different intensity averaging schemes. Two approaches were used for segmenting vasculature with the region growing tool. Firstly, by using a stringent threshold and selecting a voxel near the center of a brightly stained vessel it is possible to rapidly generate branched segmentations that do not overlap into non-vascular tissue. Secondly, individual vessels can be manually extended by selecting adjacent volumes within an overlapping sphere and careful adjustment of the thresholds for intensity values with smaller differences to non-vascular tissue. This latter mode was also used for segmenting necrotic lesions. For complex heterogeneous datasets this is needed to segment intricate structures without including adjacent voxels that represent a different tissue. To correlate with histopathology, the axes of the 3D volume were adjusted (re-registered in VGStudio Max parlance) followed by slicing through the volume to match the 2D histology image as closely as possible with three or more diseased and healthy parenchymal features (airways, blood vessels, lesions, etc.)

### Histopathology

Identification of anatomical features and pathology in the CT scans was confirmed by histological techniques using Hematoxylin and Eosin (H&E) or Masson’s trichrome stain. Briefly, samples of lung were aseptically removed and fixed in 10% buffered formalin and processed in a vacuum filtration processor using a xylene-free method with isopropanol as the main substitute fixative. Tissue sections were embedded in paraffin wax. Sections were cut at 4 µm, baked at 60°C for 15 min, dewaxed through two changes of xylene and rehydrated through descending grades of alcohol to water. These sections were stained with H&E or the Masson’s trichrome method using standard procedures. Slides were dehydrated in ascending grades of alcohol, cleared in xylene, and mounted with a mixture of distyrene, plasticizer, and xylene.

### Histology slide digitization and cross validation with µ/n-CT imaging

Human lung specimens were digitized using a Hamamatsu NDP slide scanner (Hamamatsu NanoZoomer RS2, Model C10730-12) and its viewing software (NDP.View2). The red, green, and blue color balance was kept at 100% whereas gamma correction was maintained between 0.7 and Brightness (60–110%) and contrast (100–180%) settings vary slightly between slides depending on staining quality. Resolution was ∼230 nm/pixel yielding a file size of ∼2-4.4 GB. Contrast, brightness and intensity of exported images (jpg format) were minimally adjusted using CorelDraw X8. Registration of the µ/nCT scans against histopathology images was performed manually in VGStudio Max by using blood vessels, bronchi and lesions as landmarks.

### Immunohistochemistry

Pulmonary tissue was cut into 2-4 µm thick sections, mounted on charged slides, and heated at 56 °C for 15 min. Sections were dewaxed in xylene followed by rinse in 100% ethanol and one change of SVR (95%). Slides were then washed under running water for two min followed by antigen retrieval via Heat Induced Epitope Retrieval (HIER) in Tris-sodium chloride (pH 6.0) for 30 min. Slides were cooled for 15 min and rinsed under running water for two min. Endogenous peroxide activity was blocked using 3 % hydrogen peroxide for 10 min at room temperature (RT). Slides were then washed in PBST and blocked with protein block (Novolink) for 5 min at RT. Sections were incubated with primary antibodies for cytokeratin 7 (CK7; OV-TL 12/30,DAKO,Ready-to-Use), 3-mercaptopyruvate sulfurtransferase (MPST; NBP1-82617, Novus Biologicals, 1:100), CD68 (M0814-CD68-KP1,DAKO,1:3000), LCA (M0701-CD45-2B11+PD7/26, DAKO, 1:200), MPO heavy chain (sc-34161, SantaCruz Biotechnology, 1:100), CD4 (NCL-CD4-1F6, Novocastra, 1:50), CD8 (NCL-L-CD8-295-1A5, Novacastra, 1:80), CD20 (M0755-CD20cy-L26, DAKO, 1:1000), acid sphingomyelinase (ASM; ab83354, ABCAM, 1:1000) followed by washing and incubation with either HRP anti-rabbit IgG HRP (ab6721, abcam), or the polymer (Novolink) for 30 min at RT. Slides were then washed and stained with DAB for 5 min, washed under running water and counterstained with hematoxylin for 2 min. Slides were rinsed under running water, blued in 3% ammoniated water for 30 s, washed under water, dehydrated and mounted in Distyrene Plasticiser Xylene (DPX). For isotype control sections, a similar protocol to our previous studies was followed^24,31^; either IgG4 (LS-C70325/27332) or rabbit IgG (ab37415, Abcam) was used (at the same concentration/dilution as the primary antibodies) in place of the primary antibodies (isotype control).

### IHC/Ziehl Neelsen (ZN) combination staining

The IHC protocol was followed as described above, but the hematoxylin counterstain step was eliminated, and the ZN histochemical staining was continued. Slides were incubated with heated Carbol-Fuchsin stain for 10 min and then washed under running tap water. 3% acid alcohol was applied to the slide to decolorize for 30 s or until sections appeared clear. Slides were then washed in running tap water for 2 min. Slides where then counter stained with methylene blue, rinsed under running water, dehydrated and mounted using DPX mounting media.

### Numerical analysis and plotting

Opacity plots, histograms and scatterplots were generated using Python 3.7 in the Jupyter notebook environment with the Matplotlib, Seaborn and Pandas libraries.

## Data availability

High resolution histopathology and µ/n-CT images or videos will be provided upon request or can be downloaded at: https://www.ahri.org/scientist/adrie-steyn/. Please contact Dr. Adrie J.C. Steyn

## Acknowledgements

This work was supported by NIH grants R01Al111940, R01AI134810, R01AI137043, R61/33AI138280, R21A127182, a Bill and Melinda Gates Foundation Award (OPP1130017) (AJCS) and pilot funds from the UAB CFAR, CFRB and Infectious Diseases and Global Health and Vaccines Initiative to AJCS. The research was also co-funded by CRDF Global, the South African Medical Research Council, and an NRF BRICS Multilateral grant to AJCS.

## Author contributions

Conceptualization and Design: GW, AJCS. Lung tissue preparation: GW, KN, KL. µ/nCT scanning, AP, SA. HRCT: RM, KM, LYP, MM. Pathology: PKR, TN, Histopathology: KN, KL, RLH. 3D segmentation: GW, AD, LC. Data integration: GW, JNG, AJCS. Writing initial draft: GW, JNG, AJCS. Editing: JNG, AJCS. Final draft: All authors. Figure preparation: GW, AJCS. All authors discussed the results and commented on the manuscript.

## Supplemental Figure Legends

**Figure S1.**
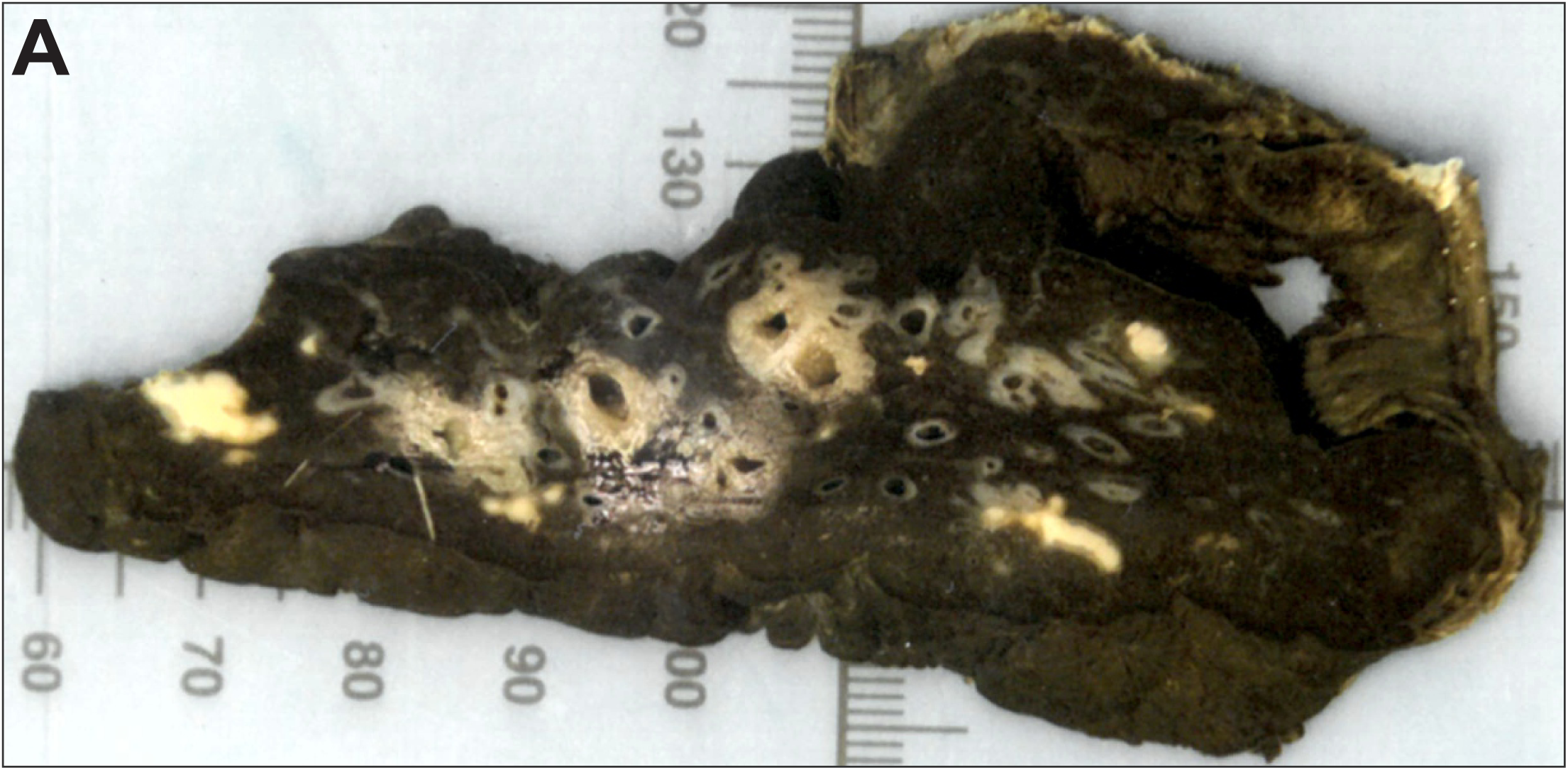
Gross image of lung section scanned using µCT. (A) Sample E; excised from a lung portion exhibiting several necrotic lesions, fibrosis, bronchiectasis, and calcification.

**Figure S2.**
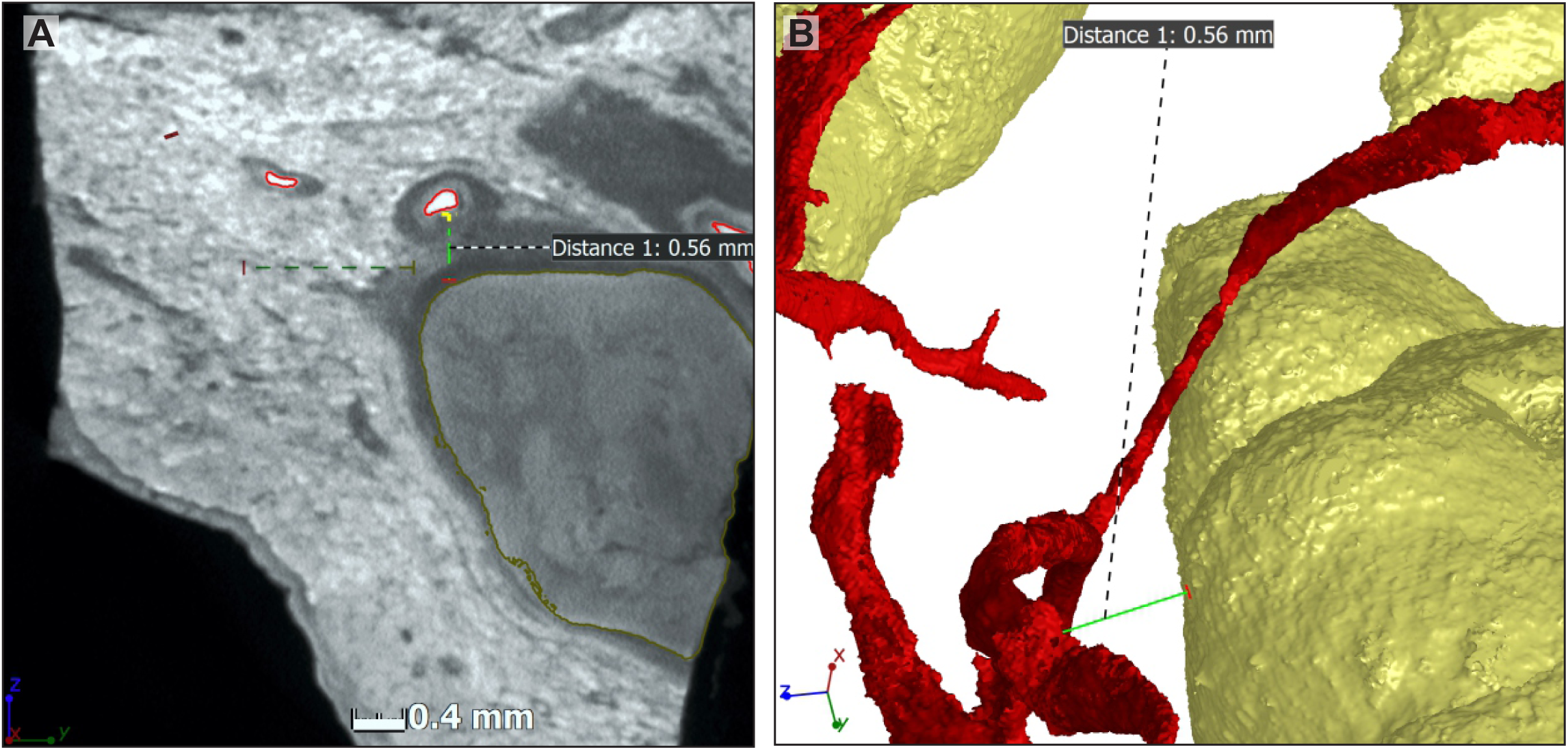
Distances between lesions and intact vasculature. Typical distances between segmented lesions (yellow) and intact vasculature (red). The µCT slice (left) and segmentation (right) are shown.

**Figure S3.**
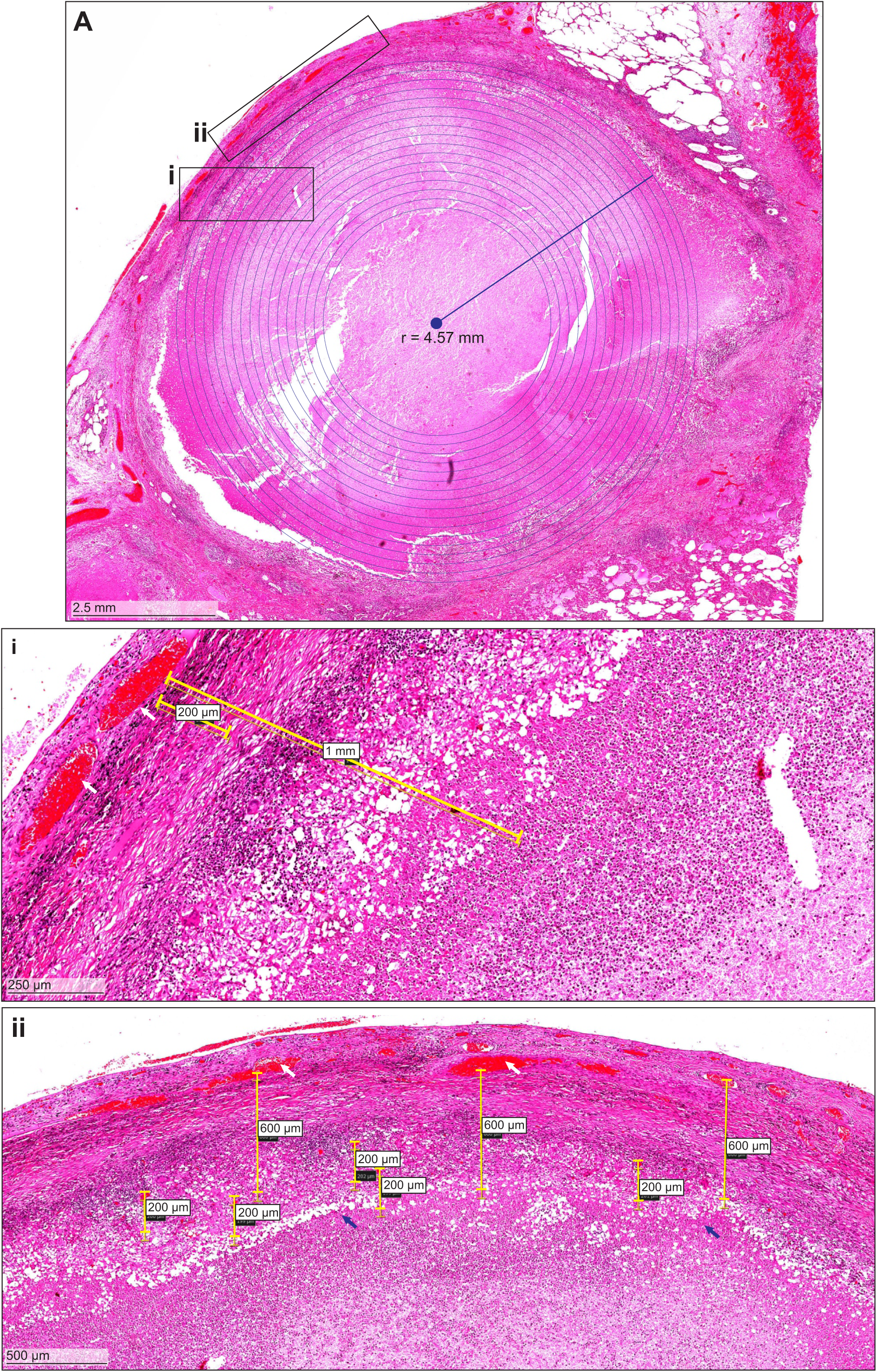
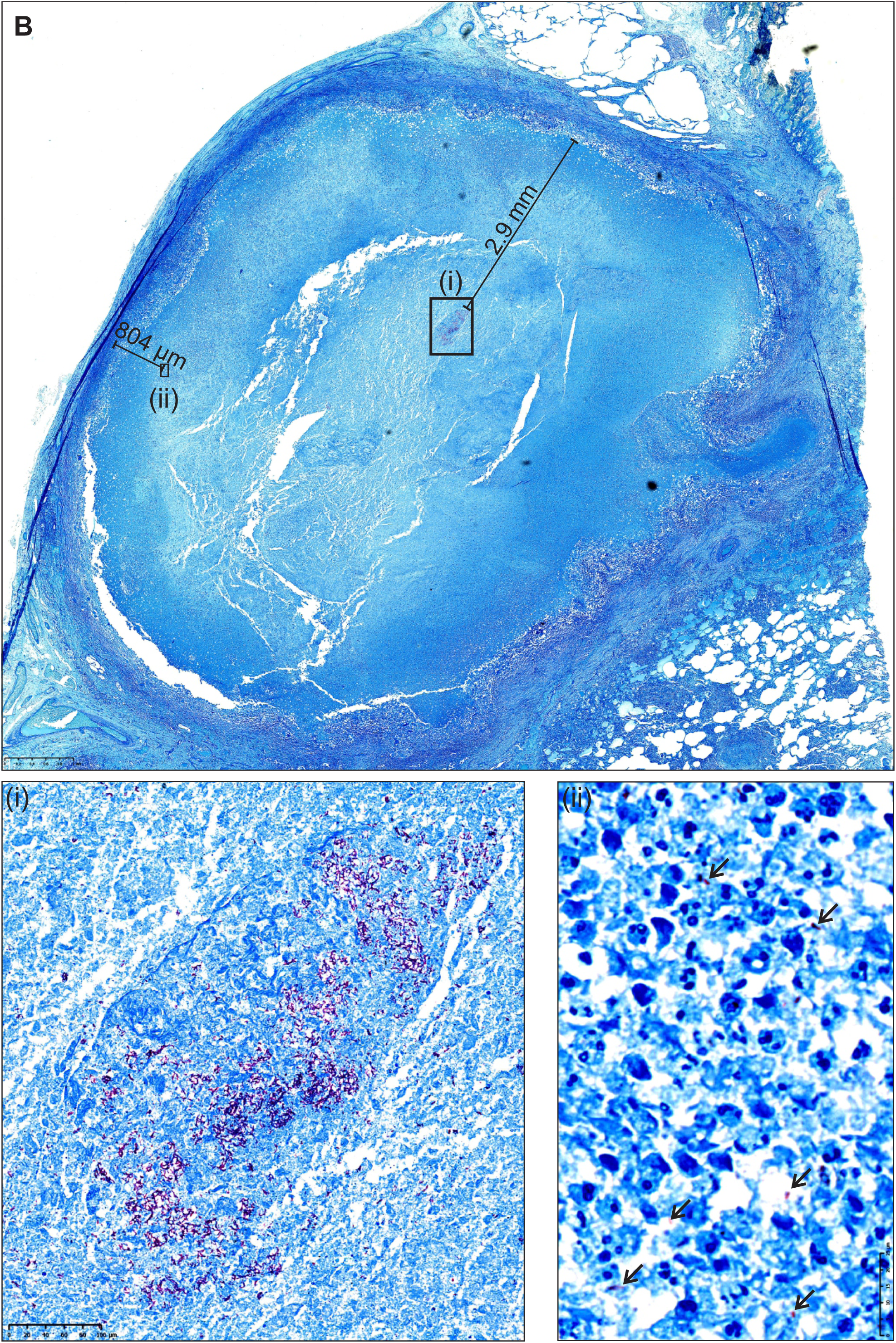
Distances between lesions and intact vasculature. H&E histology (**A**) showing distances from intact vasculature (inset i and ii) demonstrating evidence of vascular pruning of the TB lesion, strongly suggestive of lesion hypoxia. (**B**) low power depiction of ZN stain showing the distances between a large aggregate of *Mtb* cells (i), and single *Mtb* cells closest to the vasculature (ii) demonstrating that bacilli are exposed to a hypoxic environment.

**Figure S4.**
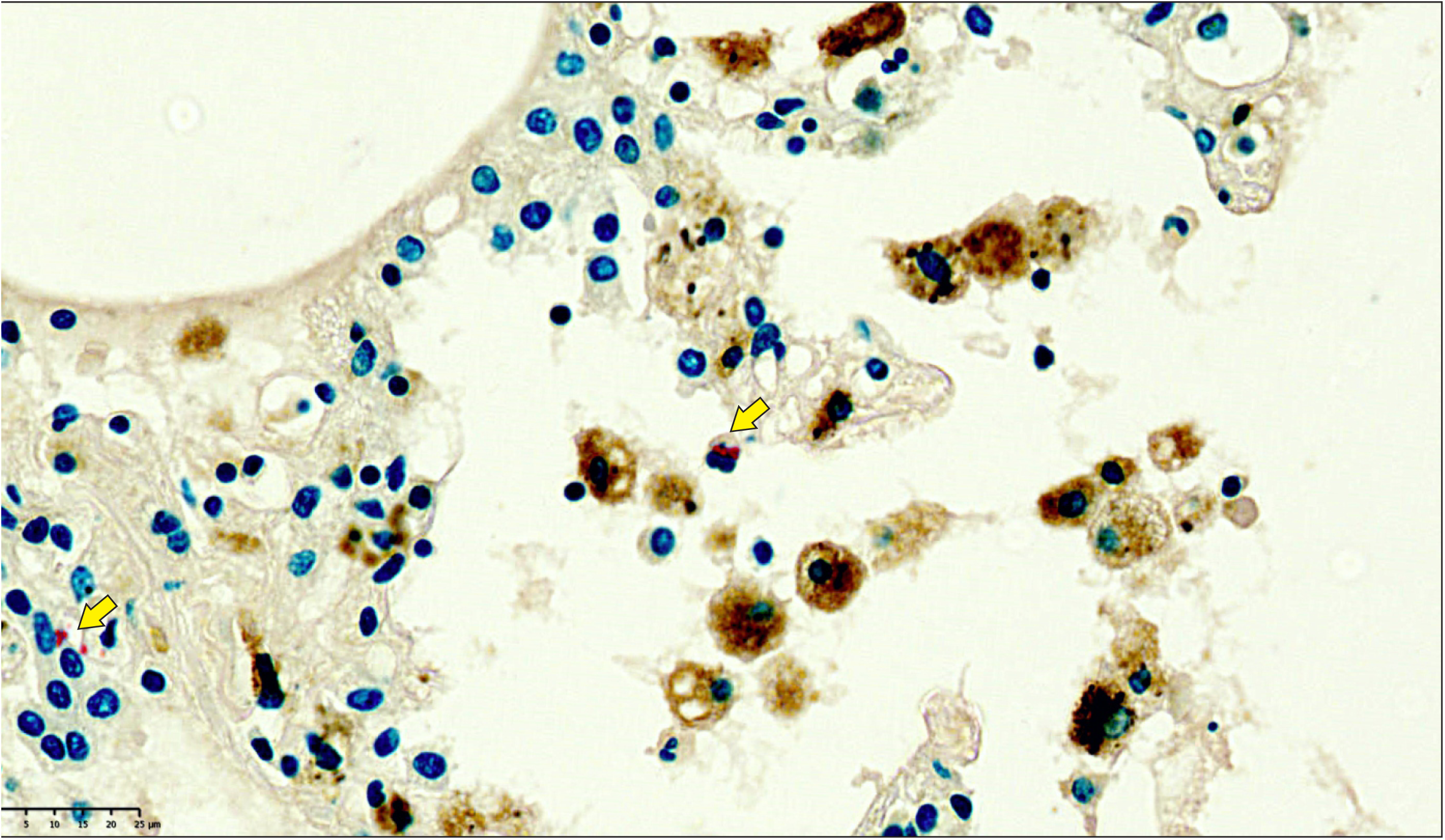
High power depiction of intracellular and extracellular *Mtb* in alveoli. Combined CD68 IHC and ZN stain. Yellow arrows indicate extracellular or intracellular (neutrophil) *Mtb*.

**Figure S5.**
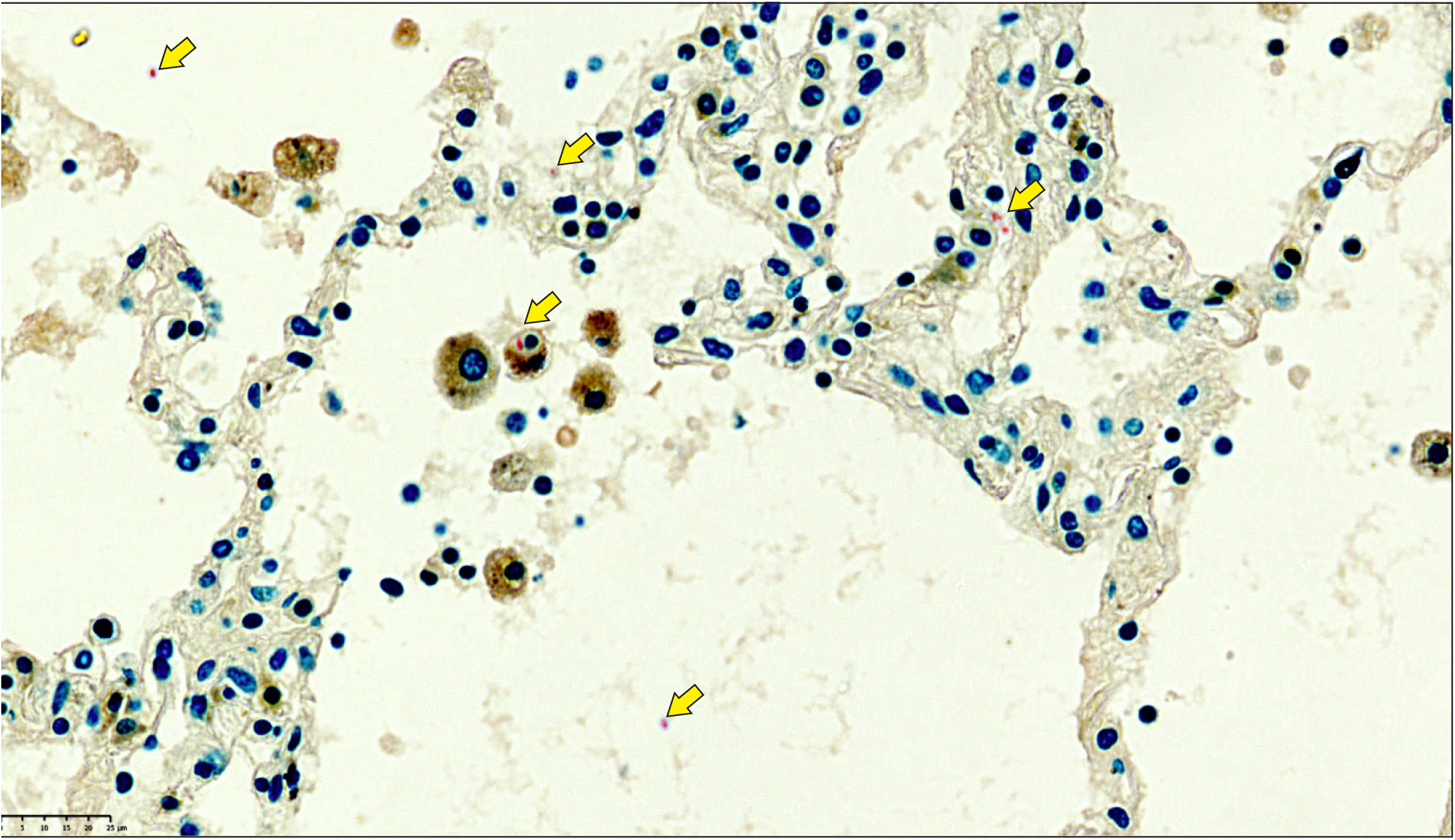
High power depiction of intracellular and extracellular *Mtb* in alveoli. Combined CD68 IHC and ZN stain. Yellow arrows indicate extracellular or intracellular (macrophage) *Mtb*.

**Figure S6.**
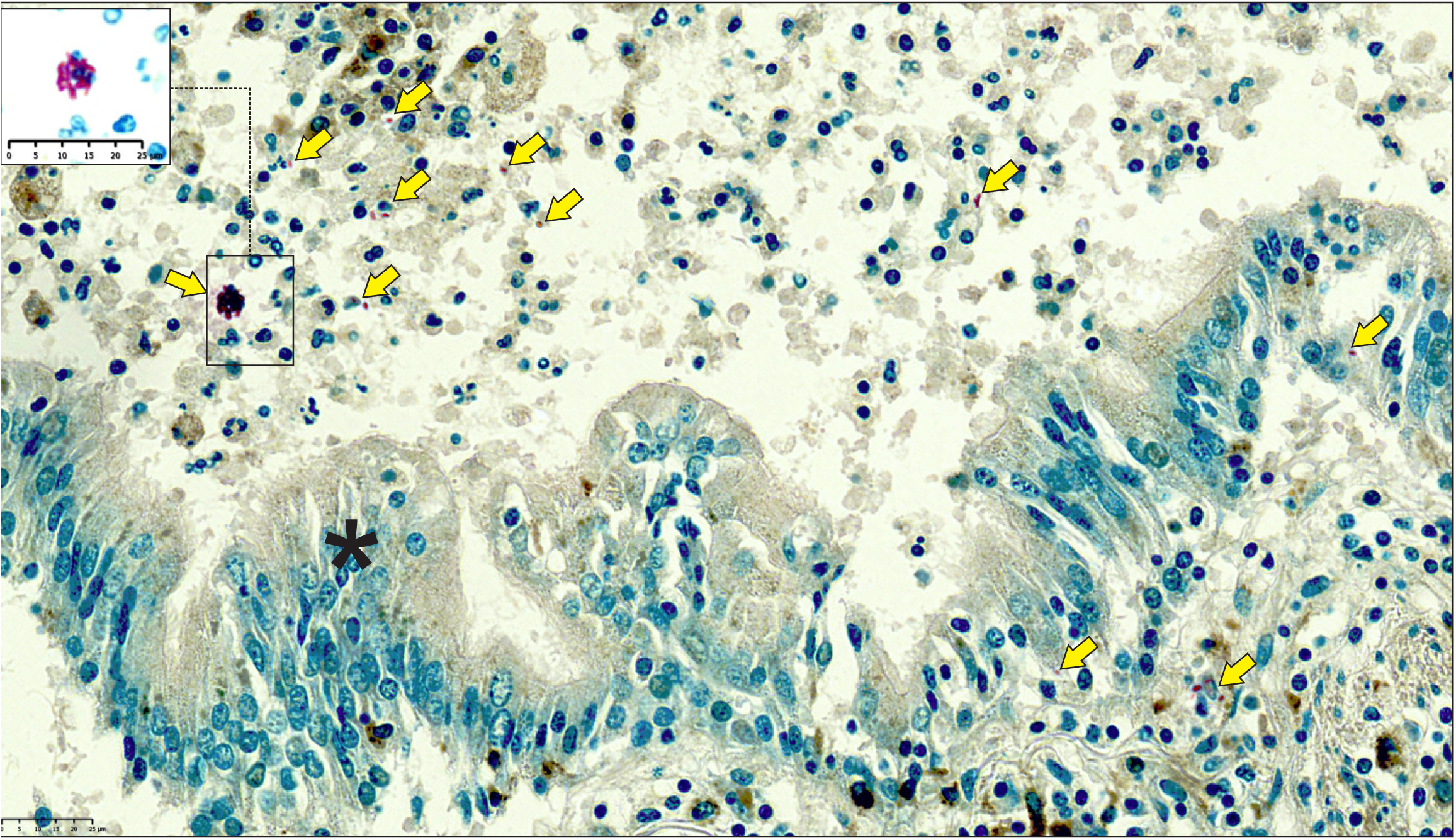
High power depiction of intracellular and extracellular *Mtb* in the luminal and adluminal areas. Combined CD68 IHC and ZN stain of an obstructed bronchus containing numerous immune cells. Yellow arrows indicate intracellular and extracellular *Mtb*. Inset; high power contrast enhanced image of a large aggregate of likely intracellular *Mtb* cells (box). *Bronchial epithelial layer.

**Figure S7.**
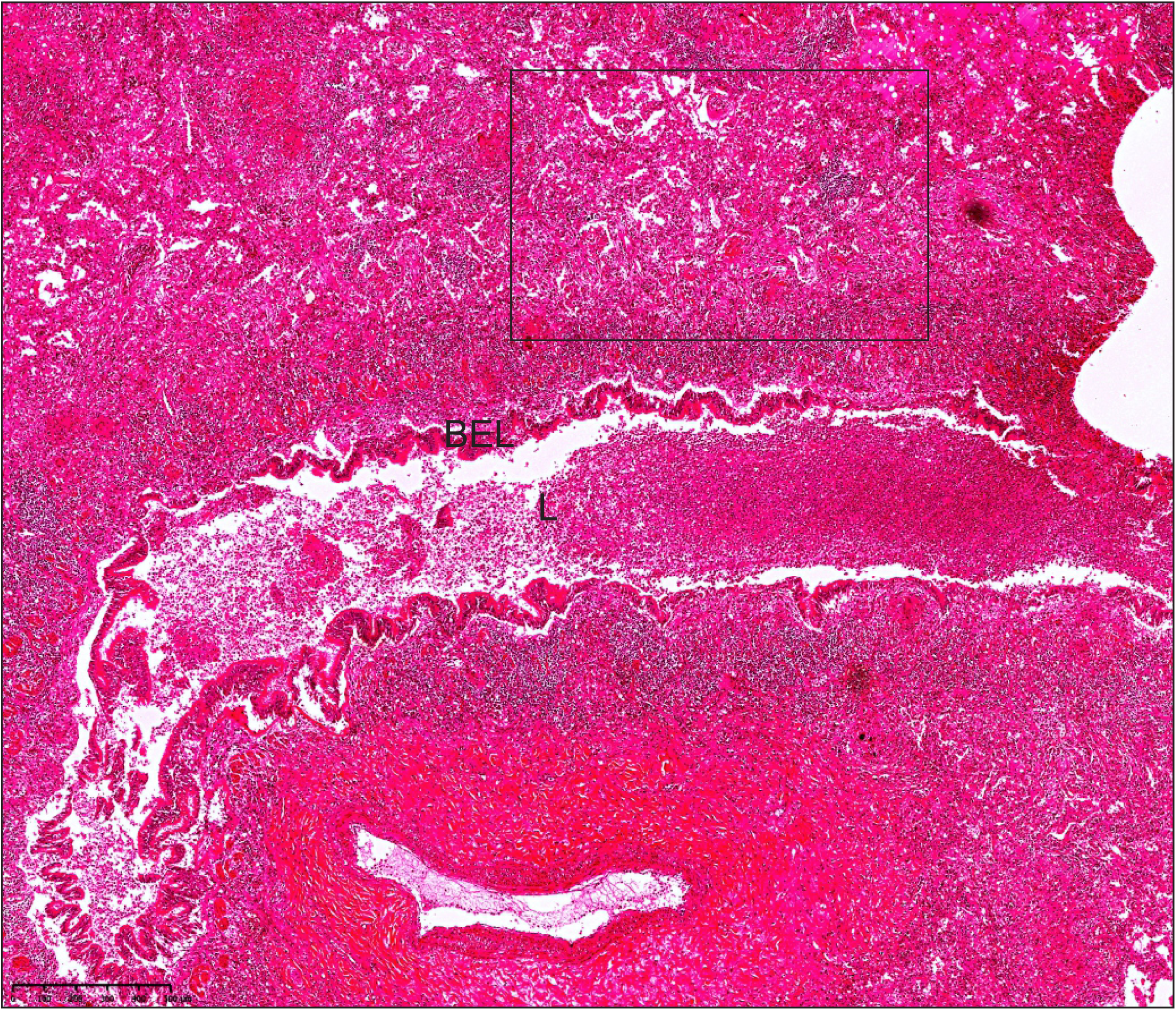
Low power image of obstructed bronchus. Low power image of H&E of obstructed bronchus used in Figure 4 (A, C, E, M, N, O, P, Q, R, S) and Figure S9. BEL; bronchial epithelial layer, L; lumen with immune cells. Box; area examined using IHC (Figure S9).

**Figure S8.**
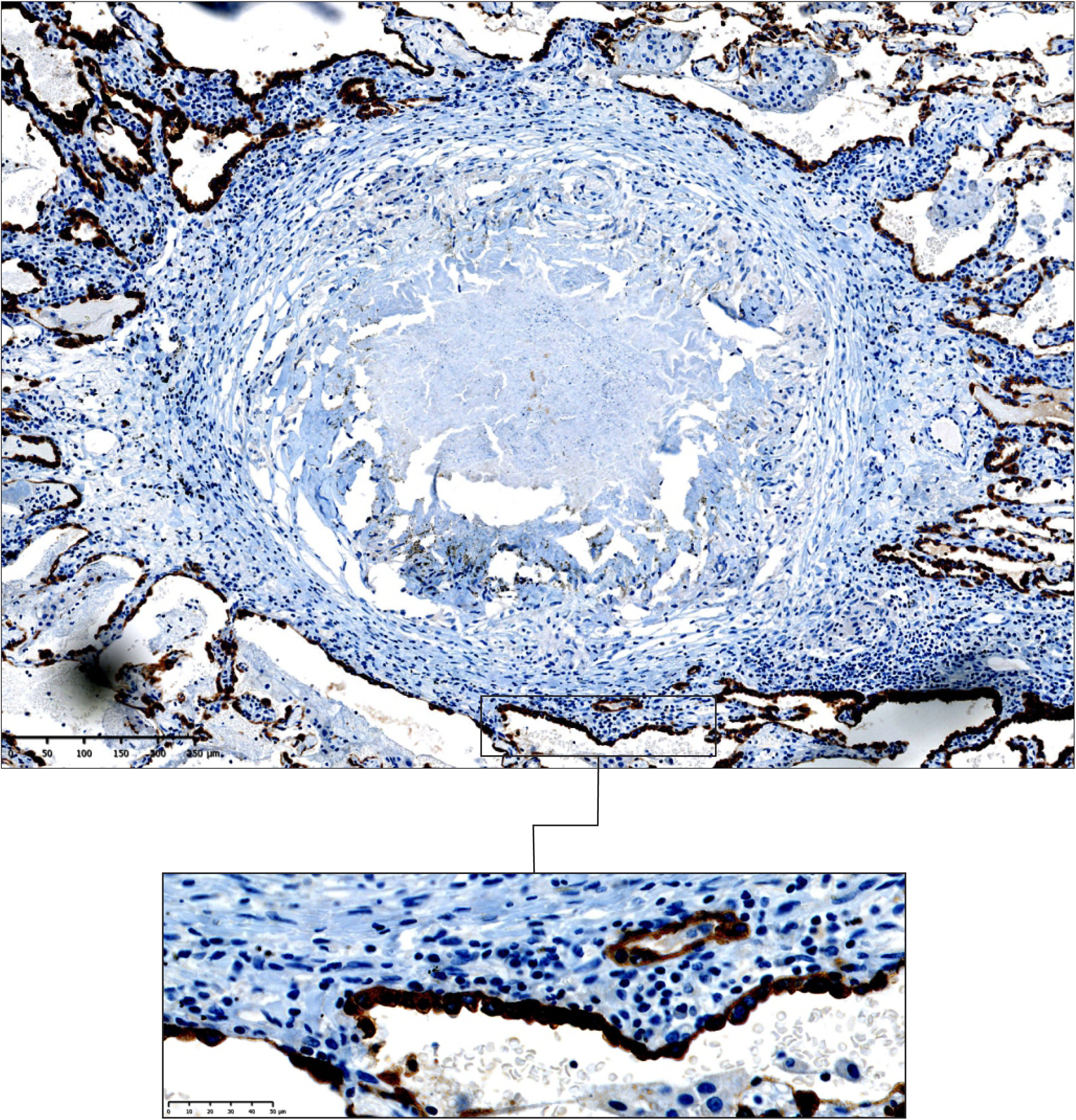
Medium power image of granuloma. Medium and high-power images of a *Mtb* granuloma stained for epithelial cells using 3MPST Ab.

**Figure S9.**
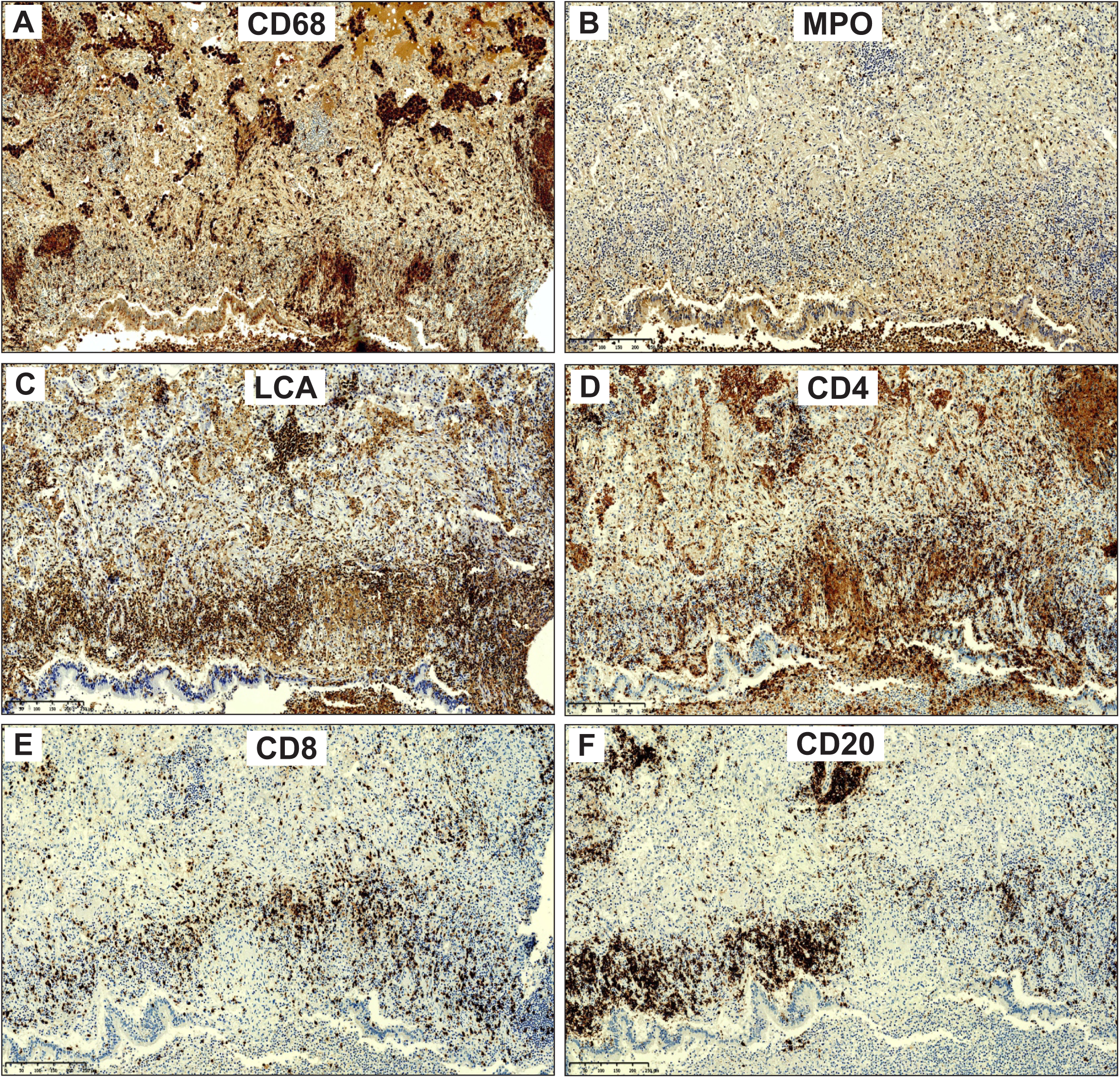
IHC of a select area in Figure 4 and Figure S7. Staining for (**A**) CD68, (**B**) MPO, (**C**) LCA, (**D**) CD4, (**E**) CD 8, and (**F**) CD20 Abs in the consolidated area of Figure S7.

**Figure S10.**
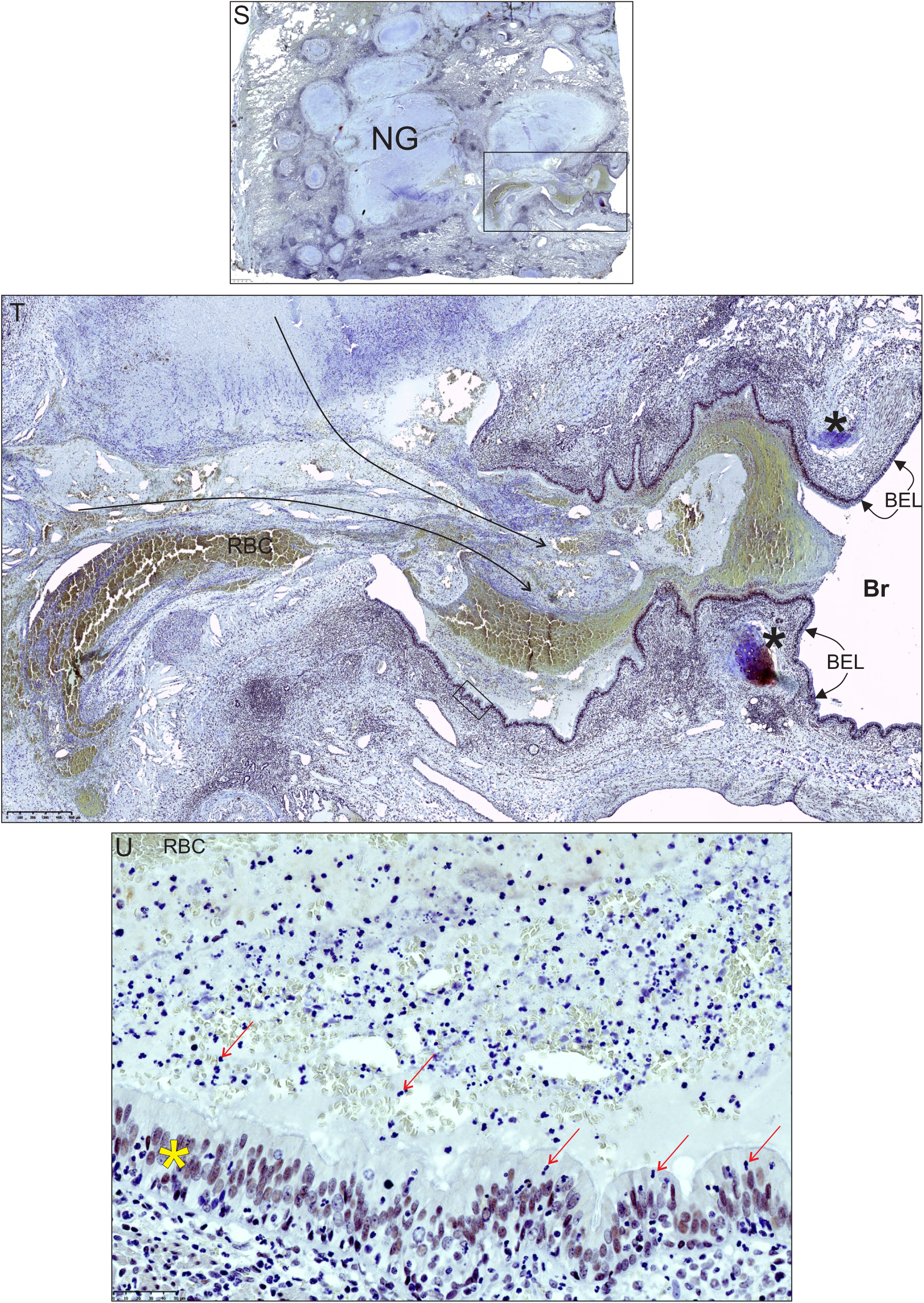
High power magnification of Figure 4 S-U.

**Figure S11.**
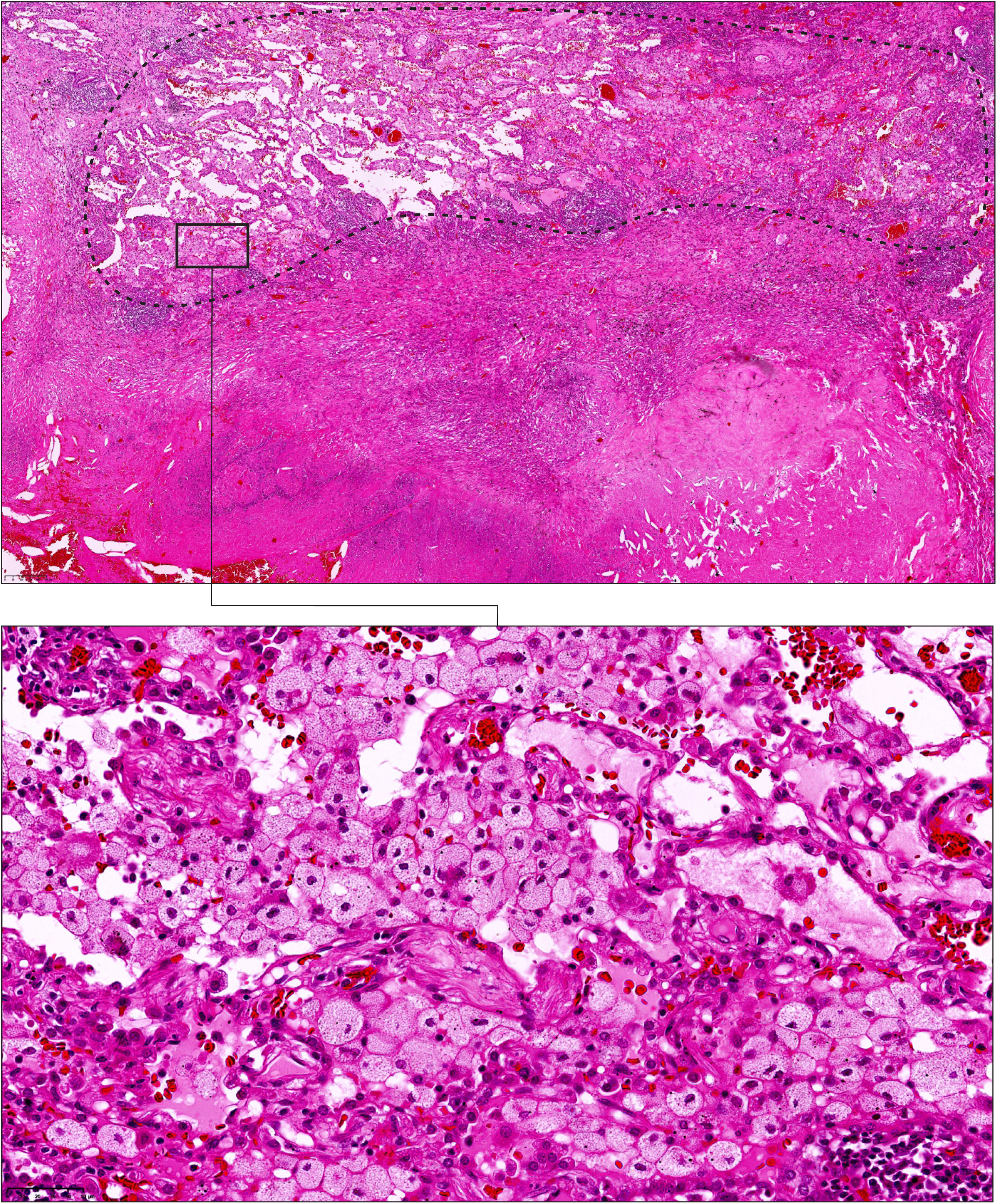
H&E stain of area containing foam cells. Low power (encircled with dotted line) and high power (boxed) images of foam cells scattered around granulomas in Figure 4 S-U and Figure S10.

## Video Legends

**Video S1. Relative orientation and directionality of caseous necrotic lesions, airways and vasculature (Sample B)**. Segmentation of caseous lesions within in slice of diseased lung reveals complex morphology, and a similar orientation to remaining airways and vasculature. Where there is a paucity of airways, lesions dominate, suggesting airway obstruction and bronchial spread of infection where the lesion acts as its source.

**Video S2. Slice video of µCT scan of caseous necrosis (Sample C)**. µCT scan through Sample M (tip of Sample B). Caseous lesions are outlined in yellow and vasculature in red. Note the cylindrical shape of the lesion. HRCT scans refer to such lesions as tree-in-bud.

**Video S3. Segmentation of caseous necrotic lesion and surrounding vasculature (Sample C, removed from Sample B)**. Caseous necrosis is outlined in yellow, vasculature in red. The vasculature “hugs” the surface of the lesions.

**Video S4. Segmentation of caseous necrotic lesion with embedded obliterated structures and surrounding vasculature (Sample D)**. Caseous necrosis is shown in yellow, vasculature in red. Obliterated structures within the lesions could also be segmented (purple), revealing a branched structure resembling a former airway.

**Video S5. Segmentation of volumes connecting necrotic lesions with airway and vascular networks (Sample A)**. Lesions (yellow), vasculature (red), airways (blue) and surrounding volumes (cyan) were segmented. The surrounding volumes around lesions, airways and vasculature are contiguous. This further demonstrate caseous necrotic lesions form within (or heavily influence) airways and are shaped by transport network morphology.

